# The fate of standing variation and new mutation under climate change

**DOI:** 10.1101/2020.01.12.903427

**Authors:** Cheng-Yueh Lu, Cheng-Ruei Lee

## Abstract

Many species face existence threat under anthropogenic climate change, and standing genetic variation was proposed as a way for sessile species to adapt to novel environments. However, it is still unknown whether standing genetic variants, being adaptive to current environmental variability, are sufficient to guarantee future survival. Here we investigate the relative importance of standing variation versus new mutations from the past to infer their future fate in nature. In the wild banana species *Musa itinerans* where the Taiwanese populations were ancient immigrants from the Chinese populations, new mutations exert larger effect size in precipitation-related variables, where Taiwan contains extreme environments beyond the ancestral climatic range, and new alleles have stronger association with novel environments. For temperature-related variables where Taiwan is within the ancestral climatic range, standing variants are more important than new mutations. The effect sizes of adaptive variants therefore differ under distinct environmental pressures, supporting theoretical predictions that natural selection favors new mutations with larger effect sizes in novel environments where the population is far from the adaptive peak. Despite their importance, large-effect variants also have higher mismatch and may be more vulnerable to future environmental perturbation, leaving minor-effect variants the main source of adaptive response to rapid anthropogenic climate change. Our work provides a support in natural environment to the previous conclusions from theoretical modeling and microbial experiments in well-controlled lab conditions.

## Introduction

Anthropogenic climatic change posts an imminent threat to most organisms. For large and sessile plant species with long generation time, the speed of migration may not keep up with environmental change, and therefore phenotypic plasticity and genetic variation in the population may allow their survival under novel environments [1,2]. Adaptive genetic variation originates from standing variants before or new mutations after environmental change. Since anthropogenic climate change greatly outpaces natural mutation, the amount of standing genetic variation is therefore critical for the rapid response of a population to changing environments [3]. It remains unclear, however, whether standing variation is sufficient to guarantee future survival, given that they were mostly adaptive to the present range of climatic variability. While it may be difficult to perform manipulative experiments in the field to compare the effects of new mutations (NM) and standing variation (SV), one could investigate NM and SV during climatic change in the past.

Adaptation could happen through genetic variants that differ in their origins (NM or SV) or effect sizes (Mendelian genes with major effects or polygenic variants with minor effects). However, how these factors interact and respond to environmental pressures remains relatively uninvestigated. For example, does SV and NM differ in their relative number or effect sizes towards environmental adaptation, and how does this relationship change with different types of environmental factor? For adaptive new mutations that were fixed when facing environment change, Fisher first predicted primarily small allelic effects [4] while Kimura emphasized intermediate effects [5]. Orr, later considering the entire adaptive walk, concluded the evolution towards a novel adaptive peak should first happen through fixation of large-effect mutations and later by small-effect polymorphisms [6]. While this was supported by some studies, the majority of these are microbial experimental evolution in well-controlled environments [7,8], and few have specifically compared the effects of NM and SV. To test whether this idea holds in nature, empirical investigations on natural populations are needed.

Taiwan is well-suited for such studies: Unlike oceanic islands such as Hawaii, Taiwan is a continental island where most species originated from the East Asian continent with recurrent gene flow [9]. The land bridge between Taiwan and China during the glacial maximum allowed exchange of SV, and the isolation during interglacial periods enabled the development of NM. Here we investigate the genomic basis of environmental adaptation of a wild banana species, *Musa itinerans*, whose habitats in Taiwan are considered peripheral from ancestral area reconstructions [10], providing an opportunity to distinguish SV from NM, as well as their response to ancestral versus novel adaptive landscapes. We investigated how past events (SV versus NM) influence present adaptation and whether local adaptation may persist under future anthropogenic climate change.

## Results

### Environmental adaptation in *Musa itinerans*

We first sampled *Musa itinerans* at 24 populations across Taiwan (Fig. 1a; fig. S1a; table S1) and investigated the population structure using 14 microsatellites (table S2). Environmental Niche Modeling (with 483 occurrence points from field survey and Google Street View) reported species distribution (Fig. 1b) in line with the previous statement that *Musa itinerans* inhabits sunny valleys, watersheds, and hillsides with gentle slopes [11]. Populations differentiated mostly between east and west (fig. S1b). The most unsuitable environments lay within the Central Mountain Range and the southwestern plains (Fig. 1b), respectively corresponding to low annual mean temperature (BIO1) and low precipitation of driest quarter (BIO17), the two most important bioclimatic variables determining species distribution (MaxEnt permutation importance [12]: 36.7 for BIO1 and 27.3 for BIO17).

**Fig. 1.**
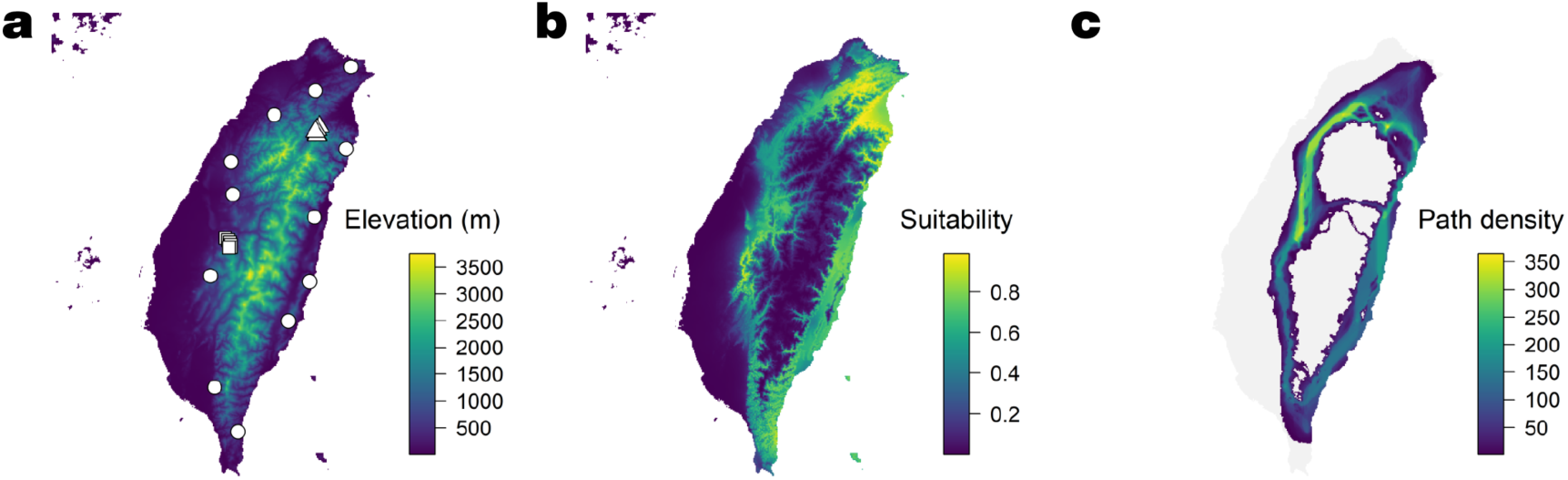
Sample distribution and niche modeling of *Musa itinerans*. **a**, Sampling sites are distributed along the latitudinal and altitudinal gradient. Lowland populations are represented as white circles; transect populations are represented as white triangles and squares. **b**, Suitability is derived from MaxEnt niche modeling. **c**, Least-cost-corridor landscape is constructed from pairwise least-cost paths among 20 populations from the resistance surface (the reciprocal of niche suitability).

To test for local adaptation, we examined the pattern of “isolation by adaptation” [13], a process where differential local adaptation restricted effective gene flow and promoted genetic differentiation among populations, by dissecting geographic and environmental effects on genetic differentiation. The strait-line “fly-over” geographical distance, calculated as the straight-line distance between locations, does not explain patterns of genetic differentiation (Mantel’s *r* = 0.146 and *P* = 0.062). However, if we considered that Central Mountain Range lacks corridors for *M. itinerans* to disperse (Fig. 1b), this fly-over geographical distance could be too unrealistic. We therefore used resistance distance, calculated from the route with least resistance among populations on the niche suitability map (Fig. 1b), to represent the “realized” geographical distance (Fig. 1c) and found that genetic differentiation was significantly associated with resistance (Mantel’s *r* = 0.226 and *P* = 0.006). The environmental Mahalanobis distance of bioclimatic variables also showed strong association with genetic differentiation (Mantel’s *r* = 0.298 and *P* = 0.005). Given that the environmental distance could be strongly dependent on geography, we performed Partial Mantel test to control the geographical effect. After controlling for realized geographic distance (resistance distance), genetic differentiation still correlates with the Mahalanobis environmental distance (Mantel’s *r* = 0.250 and *P* = 0.012), suggesting differential local adaptation is associated with genetic variation.

### Standing variation versus new mutations

To identify genomic regions associated with environmental adaptation, we performed pooled sequencing for each population. SNPs were separated into standing variation (SV: both alleles exist in Taiwan and China) or new mutations (NM, polymorphic only in Taiwan). SV outnumbered NM in both adaptive (identified with Bayenv [14]) and non-adaptive SNPs, and after controlling for the overall number of SNPs in SV and NM, SV were further enriched among adaptive polymorphisms (table S3). However, since adaptive SNPs also have higher minor allele frequency (MAF) (fig. S2a), this pattern could be confounded: SV are more likely to have higher MAF than NM, and SNPs with higher MAF may be more likely detected as adaptive due to higher statistical power. We therefore performed the same test with a subset where the adaptive and non-adaptive SNPs have similar allele frequencies (ranging from the first quantile of adaptive MAF to the third quantile of non-adaptive MAF separately for each bioclimatic variable; fig. S2a). In this case as well, SV are still disproportionately abundant (table S4), suggesting SV are more likely than NM to become environment-associated SNPs. Another potential confounding factor is the geographic extent of variants: if most NM resulted from mutations restricted to a few local populations, the limited distribution prevents environment association for NM. We therefore compared the number of Taiwanese populations containing the minor alleles for SV and NM, respectively. Contrary to the direction predicted by the aforementioned confounding factor, the geographic extent of minor alleles for SV is slightly smaller than NM (15.8 populations for SV and 16.3 for NM, *P* < 0.001).

In addition to SNP number, do NM and SV differ in their directions of effect? Under the null hypothesis that (I) the effects of NM are equally likely to facilitate adaptation to the ancestral or novel environments and (II) natural selection is equally likely to fix NM facilitating adaptation towards either direction, we expected no enrichment of new alleles in either environment. When we separated the Taiwanese populations into those within the Chinese ancestral environmental range and those with novel environments, frequencies of putatively adaptive new alleles in NM SNPs were higher in the latter set of populations, with precipitation of driest quarter (BIO17) and precipitation of coldest quarter (BIO19) showing the strongest effect (Fig. 2a; fig. S2b). Given that the directions of mutation effects should be random, these results suggest that new variants facilitating adaptation to novel environments are more likely to be retained by selection.

**Fig. 2.**
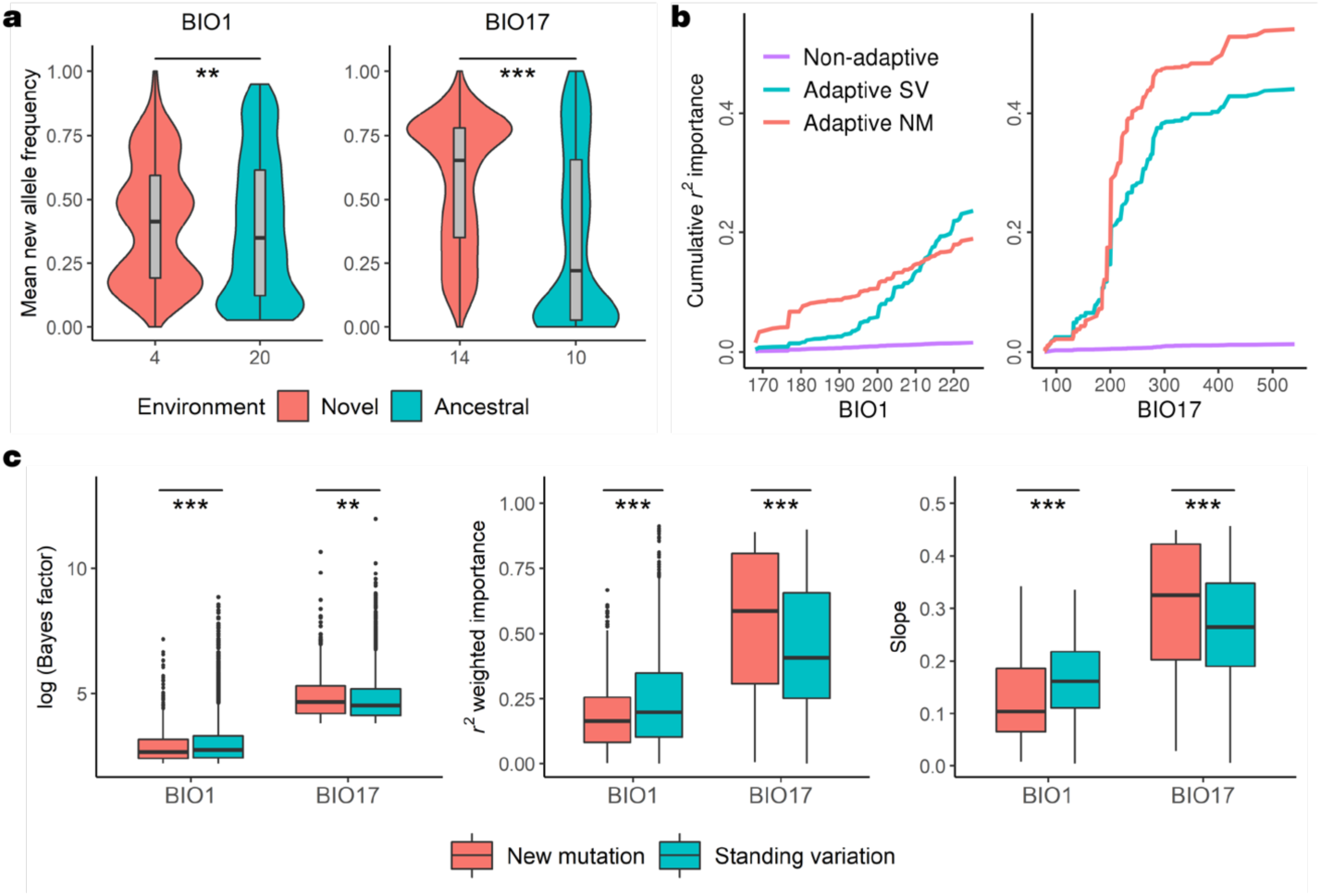
The environment-dependent enrichment of new alleles and the distinct effect sizes of standing variation (SV) and new mutations (NM) in contrasting climatic factors. BIO1 reflects annual mean temperature, and BIO17 indicates precipitation of driest quarter. **a**, Mean frequency of new alleles among NM SNPs compared between populations that have local environments within or outside of the ancestral climatic range. New alleles are enriched in novel environments (\*\**P* < 0.01, \*\*\**P* < 0.001, *t*-test). Values on the horizontal axis denote the number of Taiwanese populations within the ancestral or novel environmental range. **b**, Gradient forest cumulative *r*^*2*^ importance is shown along environmental gradients. **c**, Effect sizes as estimated from Bayes factor, gradient forest *r*^*2*^ importance, and gradient forest slope all show that SV exhibit higher effect sizes in BIO1 but the reverse in BIO17 (\*\**P* < 0.01, \*\*\**P* < 0.001, Wilcoxon rank-sum test for Bayes factor and *t*-test for *r*^*2*^ importance and slope).

The results above suggest SV might be more important than NM in terms of enriched number of variants with environment association, and adaptive NM, while lower in number, are more associated with novel environments. On the other hand, the number of candidate SNPs does not necessarily reflect the overall importance of SV over NM, since the effect size also needs to be considered. To investigate the effect size of SV and NM in environmental adaptation, we compared their Bayes factors from Bayenv and focused on two bioclimatic variables that were important in determining species distribution, annual mean temperature (BIO1) and precipitation of driest quarter (BIO17). The two variables exhibit opposite patterns (Fig. 2c): While in BIO17 and other related precipitation variables (fig. S3a) NM consistently had higher Bayes factor and therefore stronger effect size than SV, in BIO1 and other related temperature variables (fig. S3a) we observed the reverse. The same trend was observed when we estimated the effect size with gradient forest [15,16] (Fig. 2b and 2c; fig. S3b and S3c): BIO17 was the most important factor for differential local adaptation, and NM had stronger effects than SV. On the other hand, BIO1 was the least important factor where SV had stronger effects. Finally, the “importance” estimated by gradient forest is analogous to *r*^*2*^, representing the amount of allele frequency variation explained by environmental gradients. Assuming a simple linear relationship between allele frequency and environment, the value of *r*^*2*^ only represents how well each data point (a population) fits along the regression line. We were, however, also interested in the regression slope: the amount of allele frequency changes along environmental gradients (fig. S3d). Again, BIO17 had the largest overall slope among all bioclimatic variables (fig. S3e), with NM being significantly higher than SV (Fig. 2c). BIO1 had the lowest overall slope, again with the reversed pattern. Given that the MAF of adaptive NM and SV SNPs are similar, there is no need to control for allele frequency in these tests (fig. S4a). Therefore, NM with larger effect size per SNP (as estimated by Bayenv Bayes factor, gradient forest *r*^*2*^ weighted importance, and gradient forest slope) were associated with the adaptation to novel environments outside of the ancestral niche space, consistent with previous population genetics modeling results [4-6].

The observed patterns could be integrated with the unique climate of Taiwan. In comparison to the rest of the species range, northern Taiwan experiences northeastern monsoons during winter and has higher precipitation during the typical winter dry season (Fig. 3b). This pattern has been maintained since at least the last glacial maximum (Fig. 3d). The novel environments might impose novel adaptive optima to the immigrant population from China. The response to selection imposed by these environmental gradients is strong (with highest Bayes factor, *r*^*2*^, and slope among all bioclimatic variables; fig. S3a, S3c, and S3e) especially for NM, where new alleles are strongly associated with novel environments (fig. S2b). More importantly, for this major driver of adaptation (BIO17), the greatest increment of gradient forest importance lies between 200 mm and 300 mm (Fig. 2b), a range also distinguishing the novel Taiwanese versus ancestral Chinese environments (Fig. 3b). This suggests that the majority of differential local adaptation is associated with such novel-versus-ancestral environmental differences. Annual mean temperature (BIO1) is the other extreme: the environmental gradient within Taiwan is well within the ancestral Chinese environmental range (Fig. 3a), which can also be traced back to the last glacial maximum (Fig. 3c). It is likely that SV already contained genetic variants adaptive to such environmental gradients and are therefore more important than NM (Fig. 2c). In summary, we observed adaptation happening through the assortment of SV for a new territory with similar adaptive landscape and optimum (BIO1; Fig. 2c, 3a, and 3c). For adaptation to novel environments and a new adaptive landscape (BIO17; Fig. 2c, 3b, and 3d), NM with larger effect sizes were more likely favored by natural selection. Our results are therefore consistent with Orr’s model^3^, providing one of the few examples in nature.

**Fig. 3.**
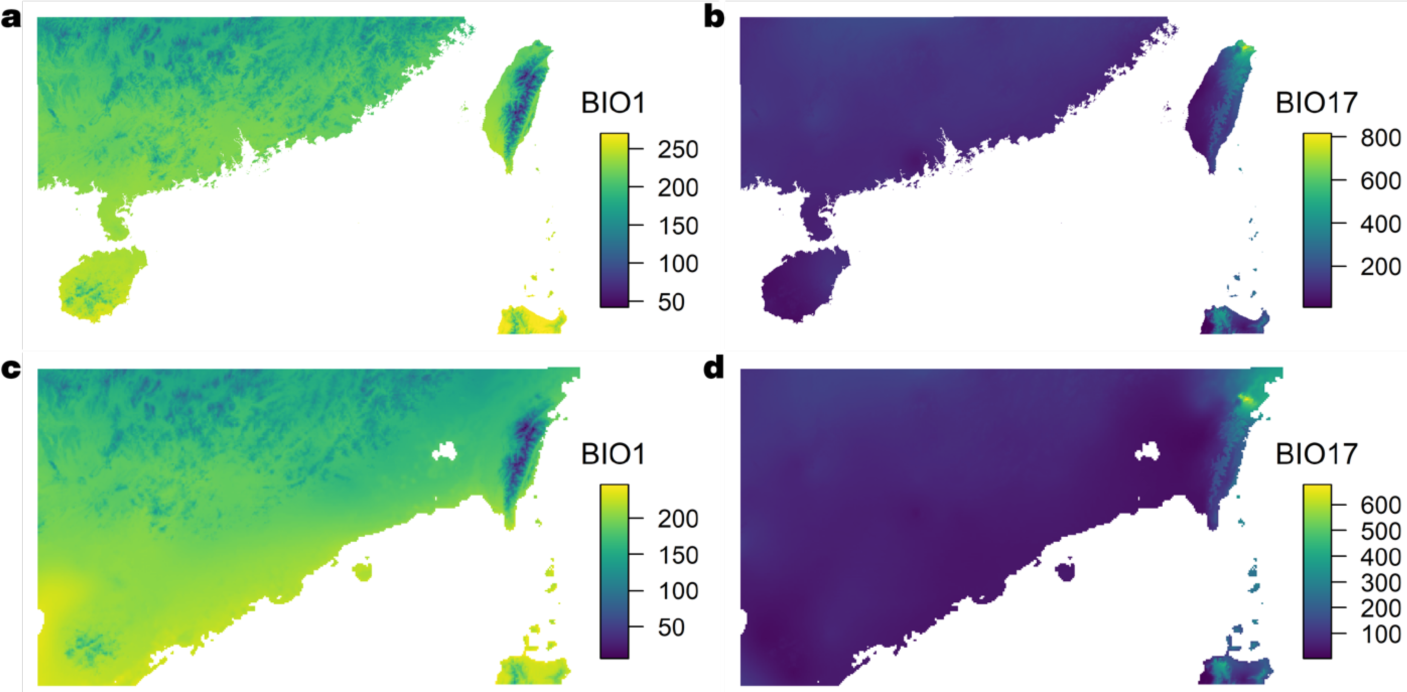
Annual mean temperature (BIO1) and precipitation of driest quarter (BIO17) for the present and last glacial maximum. **a, b**, Present environments. **c, d**, Last-glacial-maximum environments, showing the environments on the extent of land. Maps on all panels have the same range in latitude and longitude.

One key point of this study is the correct designation of SV or NM. It is possible that some SV SNPs were mis-assigned as NM if we missed an allele in China, most likely for SNPs with low MAF. We addressed this issue with the following: (I) The Taiwanese populations were nested within China in the phylogenetic tree (fig. S4b) and contained much less genetic variation (fig. S4c). Due to the stronger genetic drift in Taiwan than China, it is less likely that an originally SV SNP would retain both alleles in Taiwan but lose one in China. (II) In the extreme case, assuming 50% of NM SNPs were mis-assigned from SV, we performed 100 new analyses, each randomly assigning 50% of NM SNPs back to SV. These new analyses yielded similar results, with NM having higher effect sizes than SV in precipitation-related variables (fig. S5a). (III) Since MAF are correlated between Taiwan and China (Spearman’s rank correlation ρ = 0.35, *P* < 0.001), we performed analyses with top 50% MAF SNPs in Taiwan, thereby reducing the chance of missing minor alleles in China. The results are qualitatively the same (fig. S5b-d).

### Fate of adaptive variants under future climate change

In addition to understanding how past events (SV versus NM) affected present adaptation, we are also concerned with how these factors affect the future of this species under anthropogenic climate change. We used Bayenv to investigate the fate of currently adaptive SNPs under 16 different future climatic scenarios. Currently adaptive SNPs retaining high association with future environments are classified as “retention”, while those no longer associated with future environments are “disruption”. Different from the present pattern of SV being enriched in adaptive SNPs, we saw no clear tendency for SV or NM enriched towards retention or disruption (table S5), suggesting both types of genetic variants will be affected by climate change regardless of their previous origin.

We further used the genetic offset values from gradient forest to estimate genetic mismatch for SV and NM separately, which is associated with the magnitude of allele frequency turnover perturbed by future climatic conditions [16]. After projecting the genetic mismatch of SV and NM on the map of Taiwan for all 16 predicted future scenarios, we found SV generally has higher mean (paired t-test, *t* = 3.296, *P* = 0.005, Fig. 4A) while NM has higher maximum genetic offset across Taiwan (paired t-test, *t* = −8.608, *P* < 0.001, Fig. 4B). We further averaged the genetic offset map across all 16 future scenarios (Fig. 4C-E). There are more geographical grids with SV offset higher than NM (12,711 square kilometers) and less grids with NM offset higher than SV (8,707 square kilometers), consistent with the higher mean genetic offset for SV than NM. On the other hand, the most extreme genetic offset values were often observed for NM rather than SV (Fig. 4D-E).

**Fig. 4.**
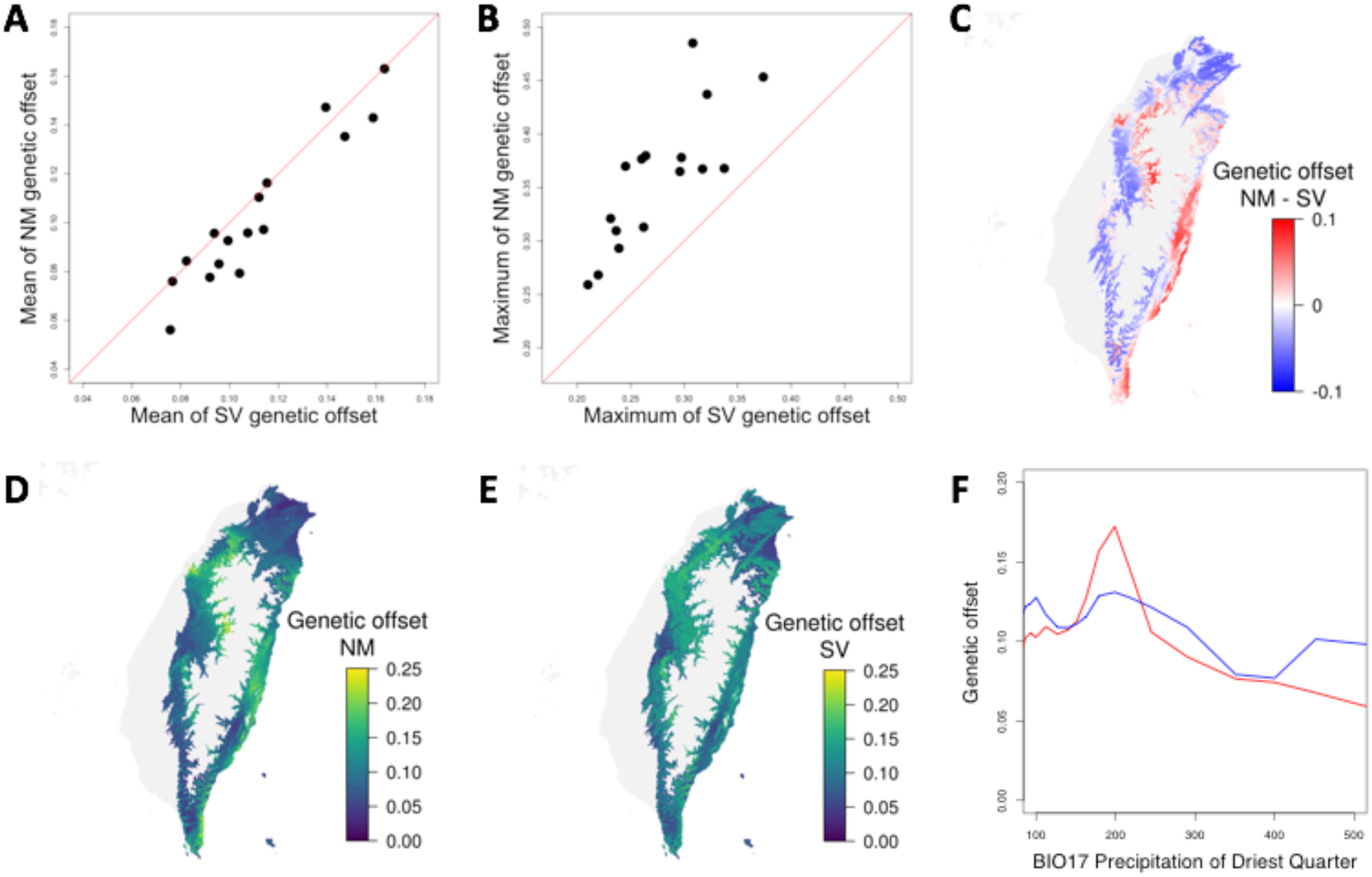
The maladaptation of standing variation (SV) and new mutations (NM) towards anthropogenic climate change. (**A**) the mean and (**B**) the maximum genetic offset of currently adaptive SV and NM SNPs across map of Taiwan for each of the 16 future climatic scenarios. In general, SV has higher mean genetic offset and NM has higher maximum genetic offset. (**C**), for the genetic offset map averaged across all 16 future scenarios, this graph shows the offset from NM minus SV. The values range from −0.07 to 0.1, representing NM has more extreme genetic offset values than SV as shown in (**D**) and (**E**). (**D** and **E**), the genetic offset value for NM and SV, averaged across all 16 future scenarios. (**F**), genetic offset values for NM (red) and SV (blue) across current gradients of BIO17 (Precipitation of Driest Quarter) across Taiwan. Grids with current niche suitability < 0.2 are excluded in all calculations and colored in gray in all maps.

Geographic regions where NM has much higher genetic offset (maladaptation towards anthropogenic climate change) than SV include northwestern mountainous and eastern coastal regions. Interestingly, these regions appear to have BIO17 values (precipitation of driest quarter) close to 200, where NM has much sharper transition of adaptive allele frequency than SV (Fig. 2B and 3B). To further examine such association, we cut the Taiwan map into 20 bins of equal grid numbers based on their current BIO17 value (excluding grids with species-wide Maxent suitability < 0.2) and estimated the mean genetic offset values of grids within each bin. Indeed, NM has much higher genetic offset in regions with BIO17 value near 200 (Fig. 4F). Such pattern exists not only for BIO17 but also for many other bioclimatic variables (fig. S6) where genetic variants (either SV or NM) tend to have higher genetic offset in environments with sharp increase in gradient forest’s cumulative importance (high delta importance per unit environmental gradient in fig. S6), indicating populations closer to the threshold of sharp adaptive allele frequency change might be more vulnerable to climate perturbation.

Taken together, for adaptation in the past, NM is relatively less enriched in number but has higher effect size per SNP. For maladaptation towards the future, here we show a similar pattern: NM has weaker genetic offset than SV in the majority of geographic regions, but NM tends to have the overall highest genetic offset values in locations near the sharp transition of gradient forest’s cumulative importance. In other words, while NM has higher effect size and sharper allele frequency transition for adaptation in the past to novel environments outside of the ancestral range, such property may also make NM more vulnerable to climate change in the future, especially near the transition zone.

## Discussion

Assessing the within-species variation in climate association is the crucial first step to understand species susceptibility to fluctuating environment, and the relative importance of standing variation (SV) and new mutations (NM) in adaptation has long been debated [17-22]. In this study, we investigate how past genetic variation (SV and NM) contribute to the adaptation of present environments as well as how these factors together affect the future fate of a species under anthropogenic climate change.

For a population facing environmental change and therefore a novel adaptive landscape, previous population genetics models have documented the effect size distribution of new mutations fixed by natural selection. Considering the entire adaptive walk of a population facing novel environments, Orr’s model predicted the distribution of effect sizes where early substitutions have larger effect than later ones [6]. Consistent with Orr’s prediction, we show that NM have stronger effect size than SV in precipitation-related variables, where Taiwan exceeds the ancestral climatic range in China (and therefore the migrating population was far away from the optimum in the new adaptive landscape). This pattern is reversed for temperature-related variables, where Taiwan has similar environmental range as China. Here we provide another perspective to recent research showing that SV contributes to adaptation [21,22]. We show that SV indeed dominate over NM in number [21]. However, the effect size of SV and NM hinges on environmental conditions: natural selection may prefer new mutations contributing to the adaptation to the new rather than the old environment, and the effect sizes of NM tend to be higher than SV under such conditions. Looking towards the future, our results imply that standing genetic variation in a population may not be sufficient for the adaptation to rapid anthropogenic climate change.

Projecting into the future, we observed similar patterns. Both SV and NM could be affected by anthropogenic climate change depending on their genetic architecture of climatic adaptation: SV has slightly larger genetic offset (maladaptation to anthropogenic climate change) than NM in the majority of geographic regions, but the largest genetic offset values were mostly observed for NM SNPs in locations near the sharp transition of gradient forest’s cumulative importance. On one hand, the higher effect sizes of NM SNPs suggest their importance in past adaptation towards novel environments. On the other, the high effect size is accompanied by sharp allele frequency transitions across environmental gradients, making NM SNPs highly sensitive (or even more vulnerable) to environmental perturbations in the future. Our results also suggest that, under frequent environmental changes such as repeated glacial cycles, although natural selection might favor new mutations with higher effect sizes under novel environments, such genetic variants might more likely be mal-adaptive during the next round of environmental change due to their high effect sizes. In other words, adaptive variants of high effect sizes may be either strongly favored or disfavored by different rounds of environmental changes, making these high-effect variants less likely to be retained as the standing variation for future. This is consistent with previous notions that genetic variants with higher effect sizes might be more likely to “overshoot” the adaptive peak [4] and may partially contribute to the observations that most adaptations have polygenic architecture [23]. The results also echo a recent experimental evolution study showing non-synonymous mutations are more difficult to be retained in antagonistic changing environments [24]. For the future fate of organisms living in natural environments, further large-effect novel mutations facilitating the adaptation to new environments may happen in the future, although they will be strongly outpaced by anthropogenic climate change.

## Methods

### Sample collection and DNA extraction

Field work was conducted during 2017 (August - December) and 2018 (January - May). We sampled *Musa itinerans* at 24 sites across Taiwan (fig. S1a; table S1). Fresh leaves were harvested from nine to fifteen individuals at each site. Total genomic DNA was extracted using the standard CTAB extraction method [25]. Since other commercial *Musa* species were also grown in Taiwan, we developed an indel marker for species delimitation. From previous studies [26-29], we identified a 6-bp insertion specific for the Taiwanese *M. itinerans* in the *atpB-rbcL* region of chloroplast. We designed a primer pair (5’-GAAGGGGTAGGATTGATTCTCA-3’; 5’-CGACTTGGCATGGCACTATT-3’) and used amplicon size to confirm all collected samples are Taiwanese *M. itinerans*.

### Simple sequence repeat genotyping and analysis

SSR primer sequences used in this study were originally developed for the genus *Musa* [30,31], which were then applied on *Musa itinerans*. Previously documented primer sequences were first searched against the *Musa acuminata* DH-Pahang genome version 2 [32] on Banana Genome Hub (https://banana-genome-hub.southgreen.fr/) to check specificity as having only one amplicon, resulting in 26 primer pairs. These primers were then experimented to check specificity on *M. itinerans*, resulting in 14 pairs (table S2). We modified each pair of primers by capping the 5’ end of forward primers with M13 sequences (CACGACGTTGTAAAACGAC) and inflorescent molecules [33]. SSR amplicons were run through capillary electrophoresis and the length of each allele was recorded.

Population structure of 20 populations (table S1) was analyzed with 14 SSR markers (table S2). Lowland populations (C35H, WFL, THNL, PTWT, P199H, MLLYT, HDPG, TTL, NAJY, HLCN, NXIR, and DFR), east transect populations (TPS300, TPS500, TPS700, and TPS900), and west transect populations (XT400, XT700, XT1200, and XT1500) were used in the analysis. We inferred the ancestry of 244 individuals with STRUCTURE 2.3.4 [34,35], parameterizing a run to have (I) run length of burnin and after-burnin period of 100,000, (II) admixture ancestry model, and (III) independent allele frequency model, further setting 20 runs for each *K* value.

To investigate the association among genetic, geographical, and environmental distance, we generated these distance matrices. Genetic distance was calculated by GenAlEx 6.503 [36,37] from 14 SSR markers; straight geographical distance (the fly-over distance) was generated by ArcGIS 10.5 (http://desktop.arcgis.com/en/); environmental distance was measured as Mahalanobis distance to address the correlation among nine bioclimatic variables (below). In addition to the fly-over geographical distance which assumes organism dispersal ignores landscapes, we further calculated as resistance distance the cumulative cost along the least cost path (below). Matrix association was examined under Mantel and Partial Mantel tests. Statistical significance was examined with 1,000 permutations. We performed Mantel tests on (I) genetic distance *vs*. fly-over distance, (II) genetic distance *vs*. resistance distance, (III) genetic distance *vs*. Mahalanobis environmental distance, and Partial Mantel tests on (IV) genetic distance *vs*. Mahalanobis environmental distance while controlling for resistance distance.

### Species distribution modeling

Current and future species distribution models were built for *Musa itinerans* using presence-only data (483 occurrence points) obtained from field survey and Google Street view. Occurrence points were then reduced to 204 cells by the removal of co-occurring presence data within the same 1×1 km grid. MaxEnt 3.4.1 [12], implemented with the maximum entropy modeling approach, reports an overall niche suitability and the importance of predictors by analyzing the presence-only data as well as background (psuedo-absence) data distribution [38-40]. We downloaded from WorldClim database version 1.4 (http://worldclim.org/) spatial layers of 19 present-day bioclimatic variables based on high-resolution monthly temperature and rainfall data [41]. Layers were selected at spatial resolution of 30 arc-second and with a mask that ranges 119.25-122.47°E and 21.76-25.49°N covering Taiwan. Variables showing high dependence (Pearson’s correlation coefficient > 0.9 calculated from ENMTools [42]) from each other were removed, resulting in nine final variables: BIO1-mean annual temperature, BIO2-mean diurnal range, BIO3-isothermality, BIO7-temperature annual range, BIO12-annual precipitation, BIO15-precipitation seasonality, BIO16-precipitation of wettest quarter, BIO17-precipitation of driest quarter, and BIO19-precipitation of coldest quarter (table S1).

Present species distribution model was constructed using the default optimization settings in MaxEnt, except the regularization set to three. We tested the predictive model by ten-fold cross-validation which was carried out by randomly partitioning the data into ten equally sized subsets and then replicating models while omitting one subset in turn. In each turn, the predictive model was built using nine subsets as training data and evaluated using the other subset as test data. The output of the predictive model is the probability of presence, or called suitability, and we averaged the ten runs to have an averaged suitability.

To predict the species distribution under different scenarios of future climate change, we projected the present-day model onto eight future climatic conditions combining two periods (2050 and 2070) and four Representative Concentration Pathways (RCP 2.6, RCP 4.5, RCP 6.0, and RCP 8.5). Future climatic layers were obtained from the WorldClim database at spatial resolution of 30 arc-second and were developed based on two general circulation models: the Community Climate System Model [43], CCSM, and the Model for Interdisciplinary Research on Climate [44], MIROC. Species distribution models for the future were carried out using the same settings described above.

To estimate the least cost path between populations, we first generated the resistance surface by taking the reciprocal of suitability. Resistance and suitability is simply a monotonic transformation in which locations with higher suitability exhibit lower resistance. Pairwise least cost path was then measured among 20 populations from the resistance surface, performed by SDM Toolbox v2.3 [45]. While least cost path is the single line with least overall cost, we also constructed the least cost corridor between populations, allowing 1%, 2%, or 5% higher cost than the least cost value. In essence, the least cost corridors represent the realized dispersal routes of organisms along suitable habitats.

### Sequencing library construction and SNP identification

We conducted whole genome pooled-sequencing [46] for each population (table S1), resulting in 24 pooled-sequencing libraries. Equal amount of DNA from ten individuals at each population were pooled, except for the PTWT population where only nine individuals were available. A library with 300-400 bp insert size for each pool was prepared using NEBNext Ultra II DNA Library Prep Kit (New England Biolabs). Libraries were then sequenced with 150 bp paired-end on the HiSeq X Ten platform.

Illumina reads were then trimmed with SolexaQA [47], followed by the removal of adaptor sequences with cutadapt [48], subsequently mapped to the *Musa itinerans* reference genome assembly ASM164941v1 [49] with BWA 0.7.15 [50]. Picard Tools (http://broadinstitute.github.io/picard) was used to mark duplicated read pairs, and the genotypes were called following GATK 3.7 best practice [51].

For the 24 pooled samples, we filtered out sites with more than two alleles, indels, quality (QUAL) < 30, quality by depth (QD) < 2, call rate < 0.74, and depth (DP) > genome-wide average depth plus three standard deviations, resulting in 4,200,177 SNPs. SNPs with minor allele frequency (MAF) < 0.05, missing data in any of the pooled-seq sample, and DP per sample < 20 were further filtered out, resulting in 1,256,894 SNPs.

To investigate the relationship between Taiwanese and Chinese *M. itinerans*, we downloaded public data from 24 Chinese accessions (SRR6382516 - SRR6382539) [52]. SNPs were called using all 24 Chinese and the 24 Taiwanese samples together following the pipeline described above. We did not perform any site filtering for this joint data set since the main objective is to investigate whether specific SNPs in Taiwan also existed in China as SV. This dataset has 18,442,853 SNPs. SRR6382532 was excluded due to high missing rate. Only when evaluating the averaged expected heterozygosity between Taiwanese and Chinese populations did we filter out sites with indels and QUAL < 30, resulting in 15,591,923 SNPs.

To assess the phylogeny of our Taiwanese populations and Chinese accessions, we downloaded *Musa acuminata* sequence (SRR7013754) as an outgroup. SNPs were called using one *M. acuminata* species, 24 Chinese and 24 Taiwanese samples together following the pipeline described above. We filtered out sites with more than two alleles, indels, QUAL < 30, and call rate < 0.9, resulting in 12,693,687 SNPs. This dataset also excluded SRR6382532.

### Environmentally-associated SNP identification

We used Bayenv 2.0 [14] to search for SNPs highly associated with environmental variables. Bayenv estimates the relationship between SNPs and environments while controlling the whole-genome population structure from a subset of loose linkage-disequilibrium SNPs. Loose linkage-disequilibrium SNPs were formed by sampling one SNP from scaffolds more than 10 kb and less than 100 kb, two SNPs from scaffolds more than 100 kb and less than 500 kb, three SNPs from scaffolds more than 500 kb and less than 1000 kb, and four SNPs from scaffolds more than 1000 kb. We then, for each bioclimatic variable, defined as the adaptive SNPs ones exhibiting top 1% Bayes factor and top 5% rho value (a nonparametric correlation coefficient capable to reduce outlier effects).

We further investigated the fate of currently adaptive SNPs under anthropogenic climate change, performing the same Bayenv analyses of currently adaptive SNPs using future climatic conditions. We included two time periods (2050 and 2070) and four Representative Concentration Pathways (RCP 2.6, RCP 4.5, RCP 6.0, and RCP 8.5) from two general circulation models, CCSM [43] and MIROC [44], resulting in 16 future climatic conditions. If a currently adaptive SNP remains strongly associated with environments, it should exhibit Bayes factor above the current threshold. We then defined as “retention” a currently adaptive SNP constantly exhibiting Bayes factor above the current adaptive threshold in all future scenarios, and defined as “disruption” a currently adaptive SNP exhibiting Bayes factor above the current adaptive threshold in none of the future scenarios.

### The gradient forest method and genetic offset

We used a novel method, gradient forest [15,16], to estimate the effect of environmental gradients on allele frequency differences among populations. Gradient forest is a regression-tree based machine-learning algorithm using environmental variables to partition SNP allele frequencies. The analysis was done separately for each SNP. The resulting “importance” measures how much of the variation in allele frequency was explained by partitioning the populations based on a specific value in an environmental variable. By making multiple regression trees (thus generating a random forest) for a SNP, the goodness-of-fit *r*^*2*^ of a random forest is measured as the proportion of variance explained by this random forest, which is then partitioned among environmental variables in proportion to their conditional importance. Such SNP-wise importance of each environmental variable is then averaged across SNPs belonging to the standing variation (SV) or new mutations (NM) group, resulting in the overall importance (of each environmental variable). In the end, one could obtain the relation curve between environmental gradient and cumulative importance (analogous to the cumulative *r*^2^, proportion of allele frequency differences among populations explained by environments). This curve has two properties. First, the highest point of the cumulative importance curve denotes the overall association between a climatic variable and allele frequency, and we used this to represent the effect size of these SNPs. Second, when traversing along the environmental gradient, a sudden increase of cumulative importance at some environmental value (for example, 20°C) denotes populations on either side of this environmental cutoff have very different allele frequency compositions. In other words, this represents a threshold factor for local adaptation.

One can use this cumulative importance curve to estimate the effect of future environmental change on local populations. In the example above, a population’s local temperature increased from 19°C to 21°C due to climate change would require larger allele frequency shift than another population whose local temperature changed from 17°C to 19°C. The “genetic offset” could then be calculated as the Euclidean distance between cumulative importance corresponding to the contemporary environmental value and that corresponding to the future environmental value, considering all bioclimatic variables together. Genetic offset can then be considered to be the magnitude of genetic change needed for a population to be still adaptive in the face of climate change.

### Regression slope

The regression slope is not given by gradient forest, since it only reports the *r*^*2*^ importance estimate. Thus, we introduced the simple linear regression *y=α+βx* to measure the regression slope. We took *y* as the allele frequency, *x* as the standardized bioclimatic variable, and *β* (slope) as the measurement of the amount of allele frequency changes along environmental gradients. By fitting simple linear regression with the general least-square approach, *β* can then be expanded to 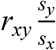, where *r*_*xy*_ is the correlation coefficient (the square root of gradient forest measured “importance”) between *x* (environment value) and *y* (allele frequency), and *s*_*x*_ and *s*_*y*_ are the standard deviation of *x* and *y*.

## Data Availability

Population pooled sequencing reads are available under NCBI BioProject PRJNA575344.

## Acknowledgements

We thank Thomas Mitchell-Olds and Sergey Nuzhdin for valuable comments, Hui-Long Chiu for the knowledge of *Musa itinerans* ecology in Taiwan, Chia-Yu Chen, Zhe-Ting Kuo, and Jo-Wei Hsieh for the assistance of sample preparation, Hao-Chih Kuo for the introduction to species distribution modeling, Cheng-Tao Lin for providing the R template of some maps, the Lee lab member for valuable discussions, and the Computer and Information Networking Center of National Taiwan University for the support of high-performance computing facilities. This work is supported by the Ministry of Science and Technology of Taiwan (106-2628-B-002-001-MY3, 107-2636-B-002-004, and 108-2636-B-002-004 to CRL).

## Author Contributions

C-RL designed the research. Both authors performed experiments and analyses and wrote the manuscript.

## Competing Interest Declaration

None

## Supplements

**Fig. S1.**
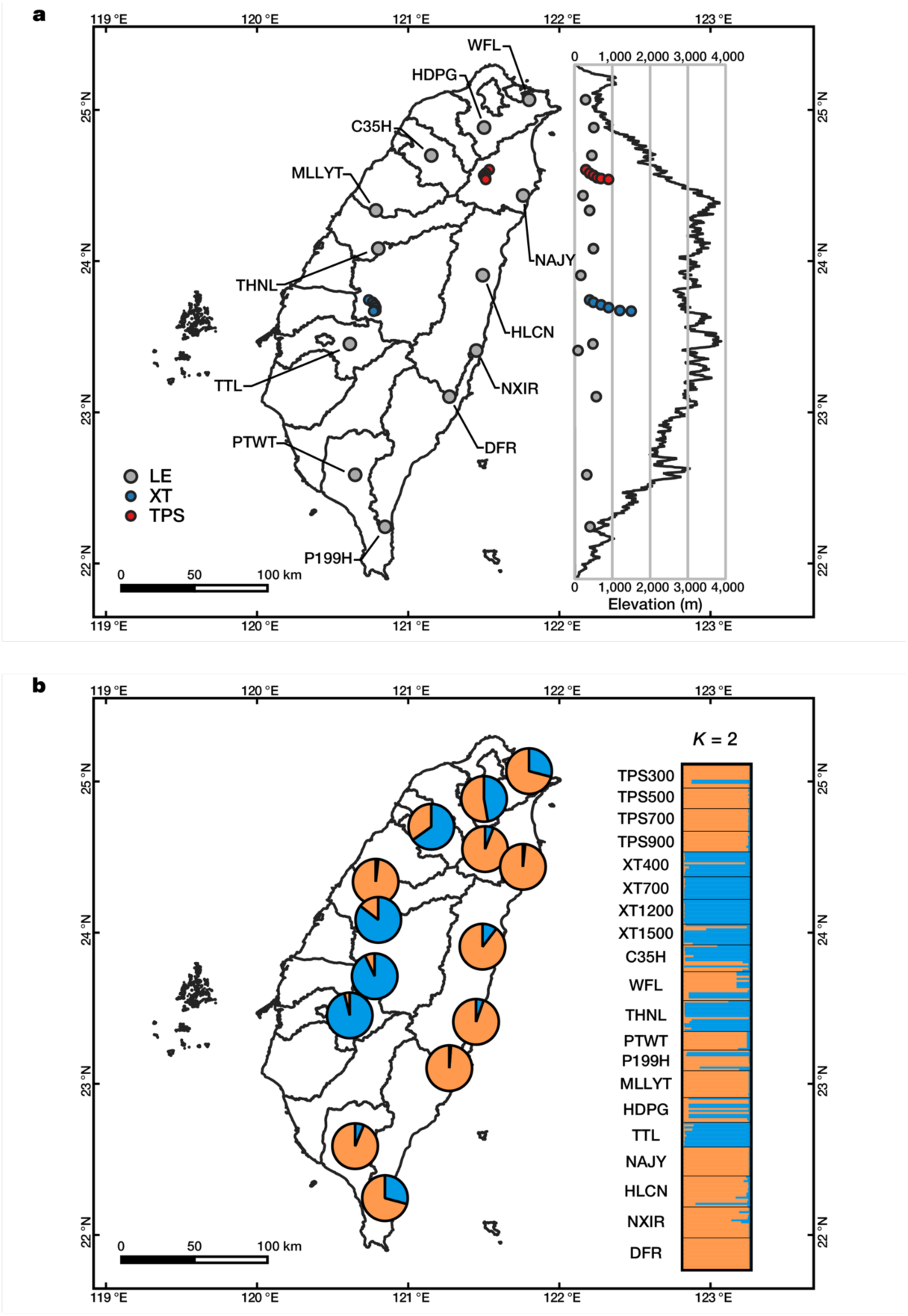
Sampling site and population structure of Taiwanese *Musa itinerans* under STRUCTURE *K* = 2. **a**, Solid circles represent collection locations corresponding to their coordinates and elevation: Gray circles indicate populations of low elevation (LE); blue circles indicate populations of Xitou transect (XT); red circles indicate populations of Taipingshan transect (TPS). **b**, Individual ancestry is plotted on the right side, while population ancestry is plotted on map with a pie chart. Map template is provided by *Cheng-Tao Lin. *Cheng-Tao Lin (2018) QGIS template for displaying species distribution by horizontal and vertical view in Taiwan. URL: https://github.com/mutolisp/distrmap_tw.qgis. DOI: 10.5281/zenodo.1493690

**Fig. S2.**
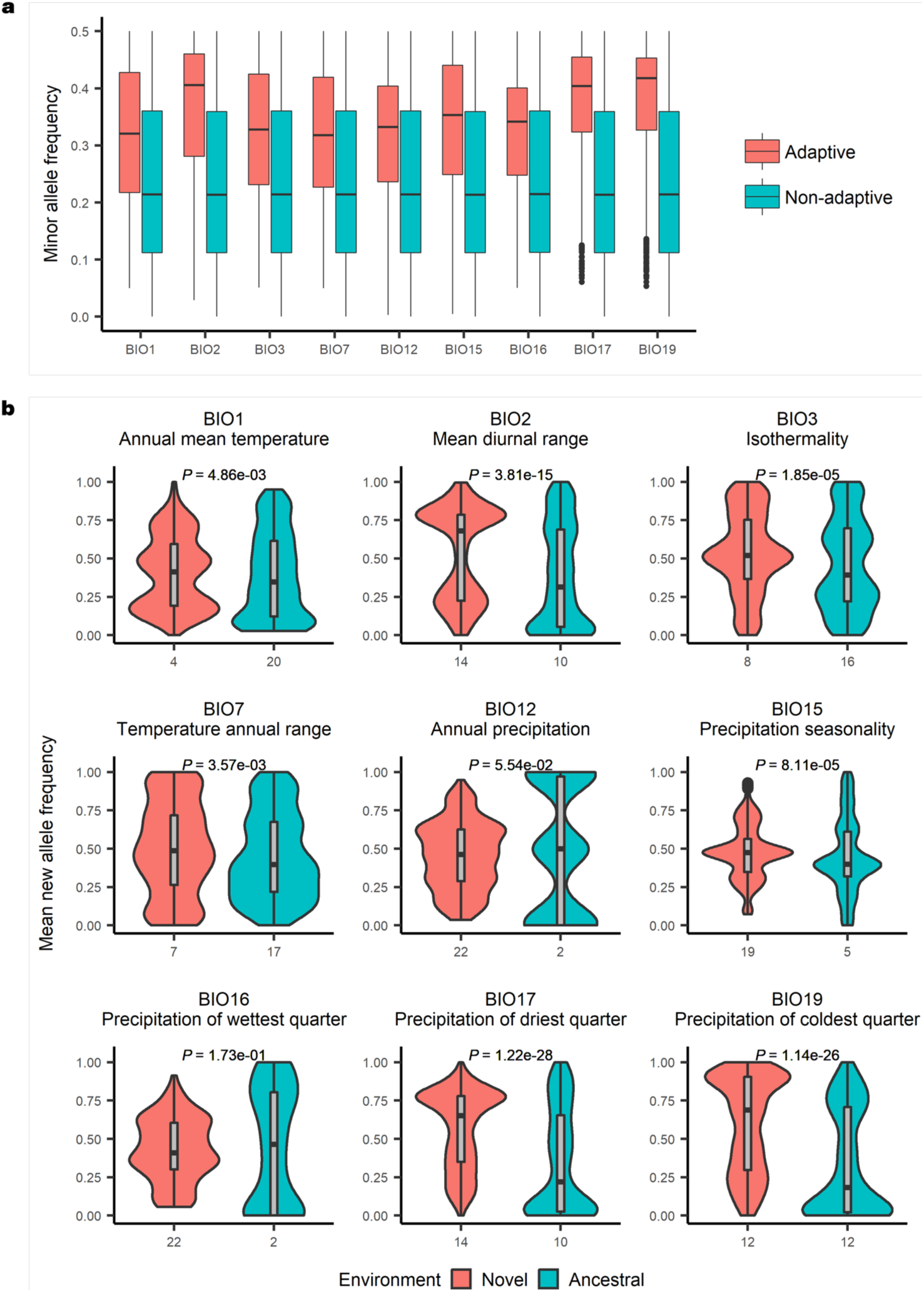
Allele frequency distribution. **a**, Minor allele frequency distribution of adaptive and non-adaptive SNPs. **b**, Frequency of new alleles in Taiwanese populations within the ancestral or novel environmental range. Statistical significance from Student’s *t*-test between the novel and ancestral environmental range for each bioclimatic variable is shown. Values on the horizontal axis denote the number of Taiwanese populations within the ancestral or novel environmental range.

**Fig. S3.**
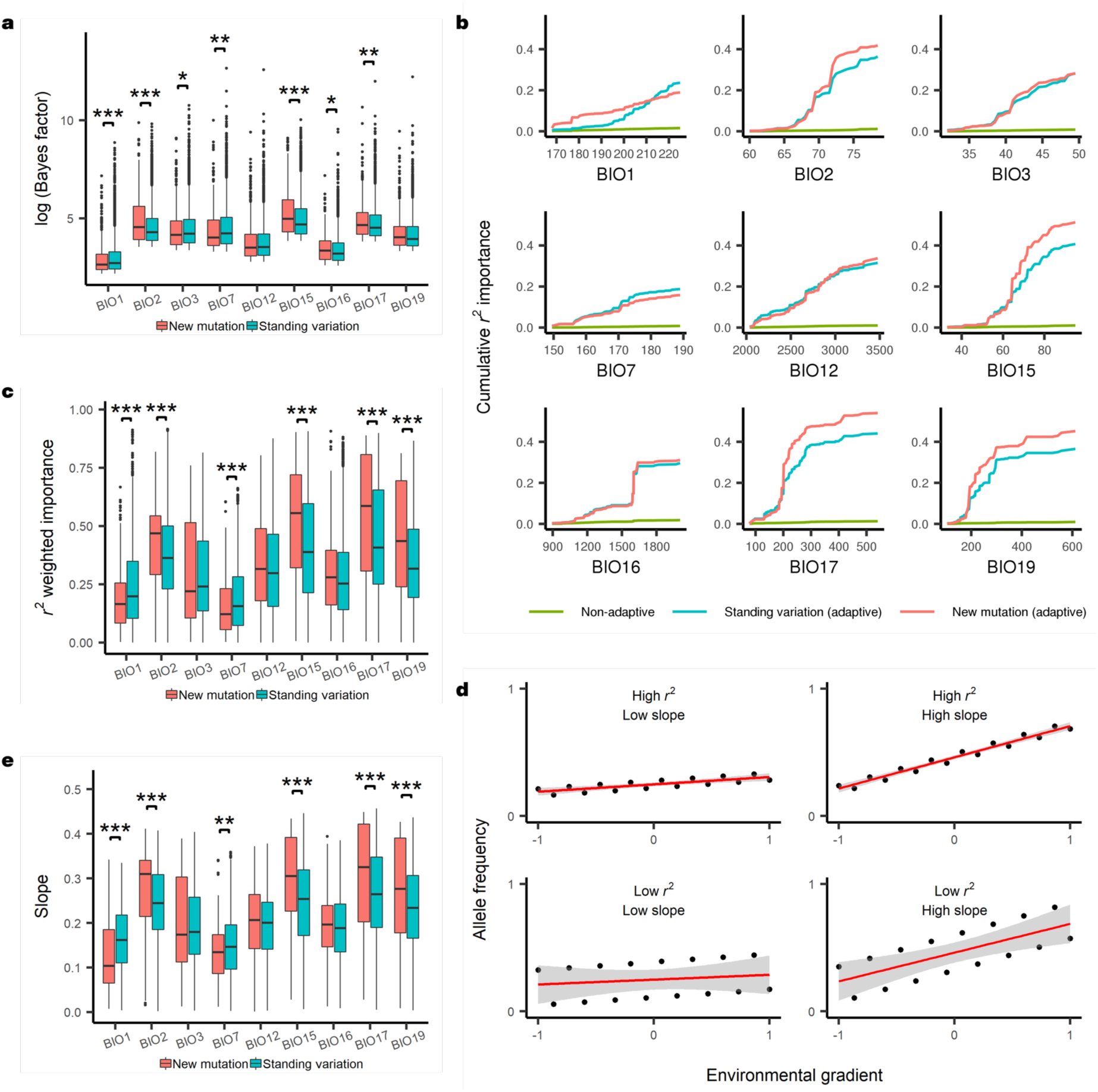
Profiling of effect sizes. **a**, Bayenv Bayes factor distribution for adaptive new mutations and standing variation (\**P* < 0.05, \*\**P* < 0.01, \*\*\**P* < 0.001, Wilcoxon rank-sum test). **b**, Cumulative *r*^*2*^ importance from gradient forest along environmental gradients. **c**, Gradient forest *r*^*2*^ importance distribution (\*\*\**P* < 0.001, Welch’s *t*-test). **d**, Example figure showing the relationship between *r*^*2*^ importance and slope. *r*^*2*^ indicates the extent that allele frequency fits a linear model, while slope indicates the amount of allele frequency changes along the linear relationship. The graphs indicate one should also investigate regression slopes in addition to the gradient forest *r*^*2*^. Values on the horizontal axis show the range of standardized environmental variables. **e**, The distribution of regression slopes when one regresses adaptive SNP allele frequency onto environmental gradients (\*\**P* < 0.01, \*\*\**P* < 0.001, Welch’s *t*-test).

**Fig. S4.**
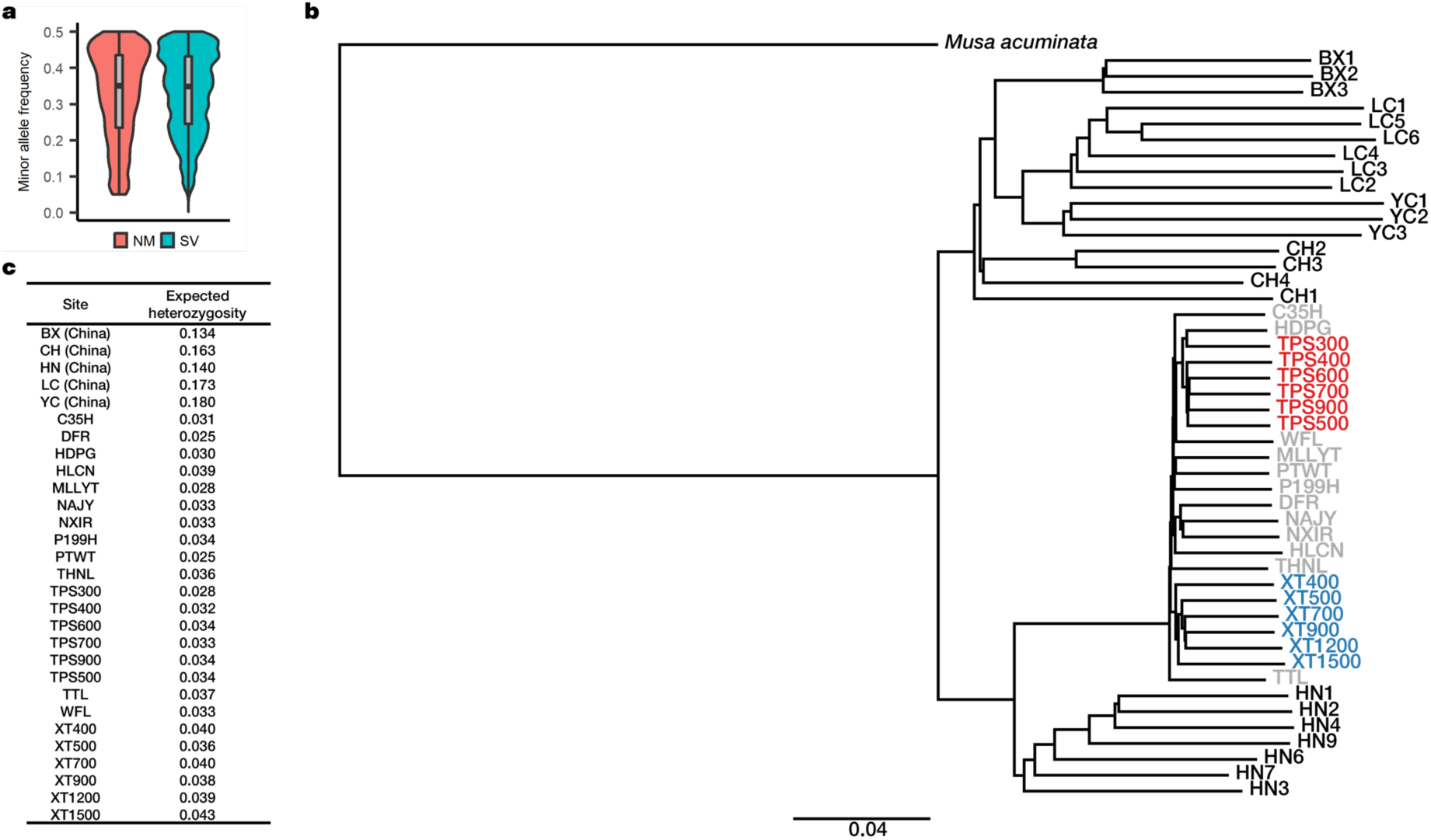
Evolutionary analyses on Taiwanese and Chinese *Musa itinerans*. **a**, Minor allele frequency distribution of adaptive SNPs in Taiwan. **b**, Phylogeny of Taiwanese and Chinese *Musa itinerans*. The gray-colored indicates Taiwanese lowland populations (LE); the blue-colored indicates Taiwanese populations of Xitou transect (XT); the red-colored indicates Taiwanese populations of Taipingshan transect (TPS); the black-colored indicates Chinese accessions and the outgroup *Musa acuminata*. **c**, Table showing the expected heterozygosity among Taiwanese and Chinese populations.

**Fig. S5.**
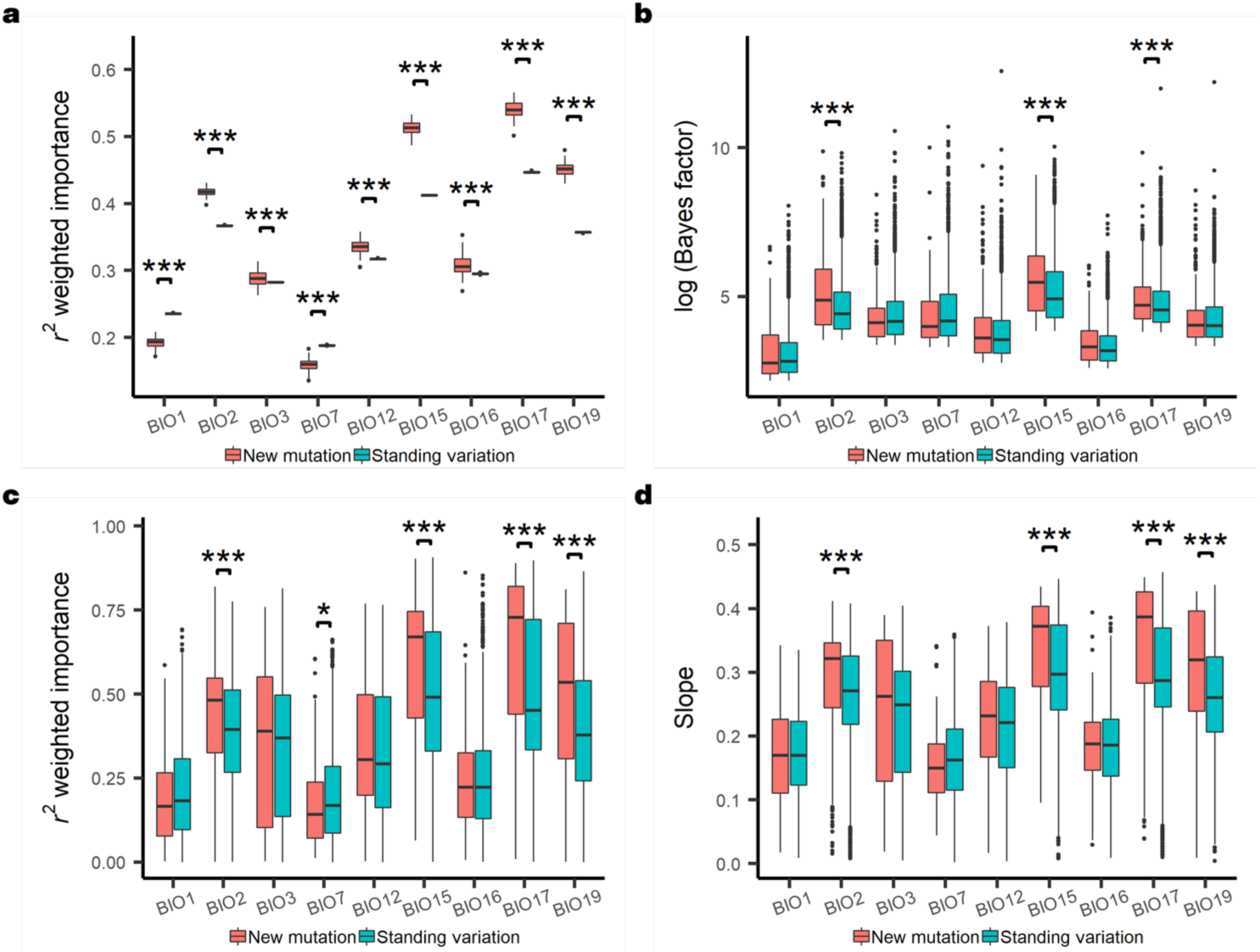
Multi-analyses addressing ascertainment of standing variation and new mutation. **a**, The distribution of gradient forest *r*^*2*^ importance across 100 re-sampling trials. In each re-sampling trials, a random set of 50% new mutations were assigned as standing variation, and the mean *r*^*2*^ importance was reported for each trial. All comparisons show strong statistical significance from Student’s *t*-test between new mutations and introduced standing variation (\*\*\**P* < 0.001). **b**-**d**, Profiling of effect sizes of adaptive SNPs with top 50% minor allele frequency. Distribution of Bayes factor (\*\*\**P* < 0.001, Wilcoxon rank-sum test; **b**), gradient forest *r*^*2*^ importance (\**P* < 0.05, \*\*\**P* < 0.001, Welch’s *t*-test; **c**), and regression slopes of adaptive SNP allele frequency onto environmental gradients (\*\*\**P* < 0.001, Welch’s *t*-test; **d**) is analyzed between new mutations and standing variation for each bioclimatic variable.

**Fig. S6.**
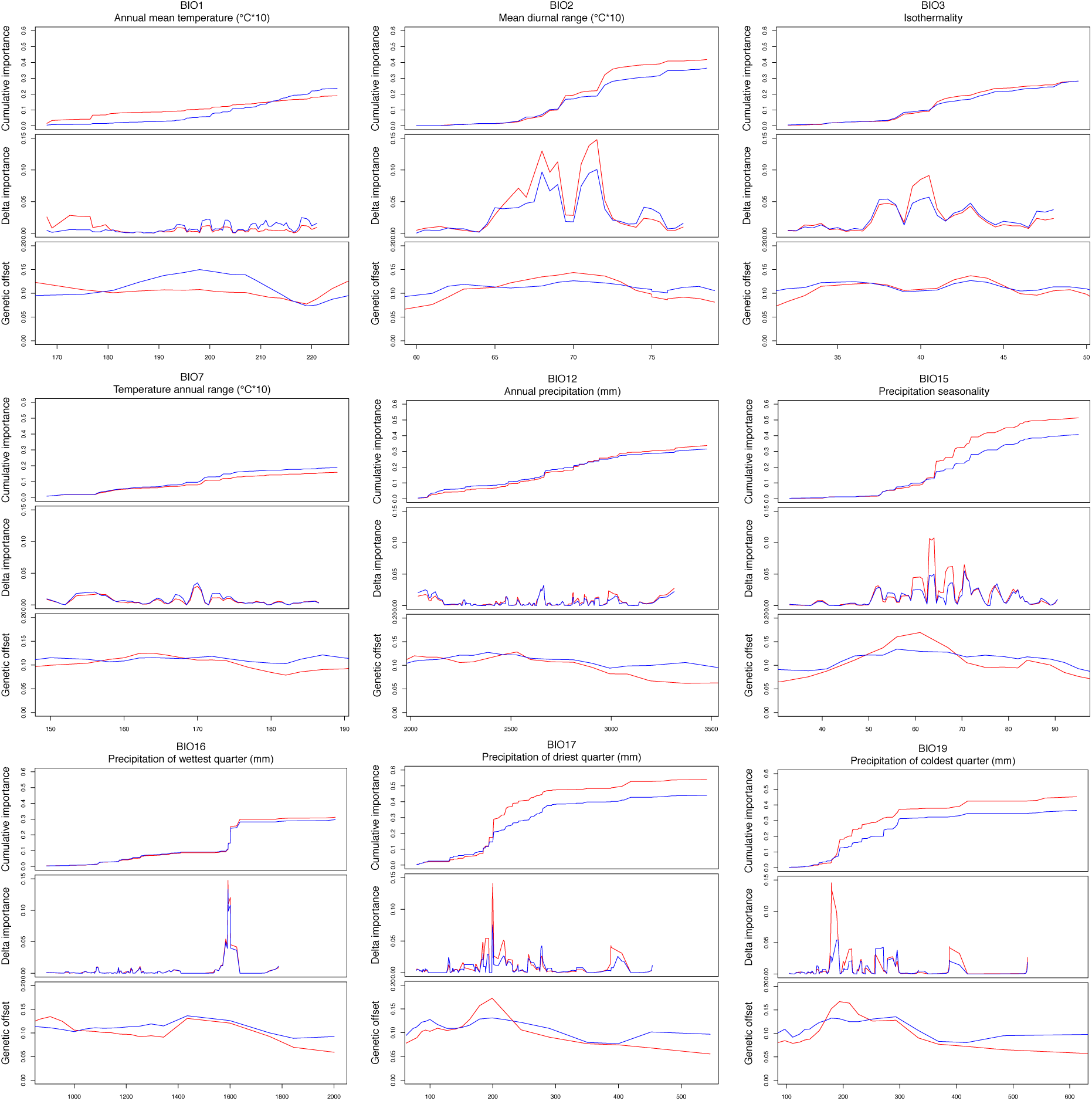
Association among gradient forest cumulative importance, delta importance per unit environmental gradient, and genetic offset for each bioclimatic variable. In each figure, red line represents new mutation (NM), and blue line represents standing variation (SV). The “delta importance” is analogous to the instantaneous slope on the cumulative importance graph. Genetic offset was calculated for SV and NM separately, where the genetic offset maps of 16 future climatic scenarios were averaged into one map for SV and one for NM. Geographic grids were separated into 20 bins based on their bioclimatic values, and the mean genetic offset in each bin was plotted. Grids with current niche suitability < 0.2 are excluded in all calculations and colored in gray in all maps.

**Table S1.** Population coordinates and bioclimatic information.

| Site | Latitude | Longitude | BIO1 | BIO2 | BIO3 | BIO7 | BIO12 | BIO15 | BIO16 | BIO17 | BIO19 |
| --- | --- | --- | --- | --- | --- | --- | --- | --- | --- | --- | --- |
| TPS300 | 24.60488 | 121.5359 | 208 | 63 | 35 | 180 | 2,970 | 46 | 1,223 | 364 | 364 |
| TPS400 | 24.58135 | 121.5124 | 194 | 65 | 36 | 177 | 2,769 | 50 | 1,145 | 301 | 301 |
| TPS500 | 24.56765 | 121.4992 | 199 | 65 | 36 | 179 | 2,763 | 50 | 1,142 | 299 | 299 |
| TPS600 | 24.55172 | 121.5090 | 193 | 65 | 36 | 176 | 2,709 | 51 | 1,124 | 289 | 289 |
| TPS700 | 24.54638 | 121.5125 | 185 | 65 | 37 | 174 | 2,699 | 49 | 1,107 | 302 | 302 |
| TPS900 | 24.54021 | 121.5126 | 178 | 66 | 38 | 173 | 2,757 | 46 | 1,110 | 336 | 336 |
| XT400 | 23.74292 | 120.7386 | 217 | 77 | 42 | 180 | 2,468 | 86 | 1,395 | 87 | 127 |
| XT500 | 23.72780 | 120.7641 | 206 | 77 | 44 | 173 | 2,600 | 84 | 1,420 | 97 | 141 |
| XT700 | 23.71137 | 120.7782 | 199 | 77 | 45 | 169 | 2,412 | 83 | 1,311 | 94 | 138 |
| XT900 | 23.69294 | 120.7857 | 189 | 77 | 47 | 162 | 2,357 | 83 | 1,287 | 97 | 145 |
| XT1200 | 23.67333 | 120.7854 | 176 | 77 | 48 | 159 | 2,478 | 82 | 1,346 | 97 | 148 |
| XT1500 | 23.66936 | 120.7710 | 160 | 78 | 48 | 160 | 2,637 | 81 | 1,402 | 98 | 149 |
| C35H | 24.69907 | 121.1528 | 202 | 65 | 34 | 188 | 2,500 | 55 | 1,073 | 192 | 289 |
| WFL | 25.06752 | 121.7998 | 203 | 58 | 31 | 185 | 3,440 | 29 | 1,229 | 606 | 749 |
| THNL | 24.08273 | 120.8021 | 212 | 74 | 40 | 184 | 2,188 | 83 | 1,220 | 78 | 138 |
| PTWT | 22.58684 | 120.6489 | 227 | 78 | 49 | 158 | 2,912 | 98 | 1,784 | 79 | 87 |
| P199H | 22.24302 | 120.8465 | 223 | 73 | 50 | 144 | 3,114 | 92 | 1,856 | 178 | 178 |
| MLLYT | 24.33526 | 120.7864 | 211 | 71 | 37 | 190 | 2,056 | 76 | 1,084 | 79 | 148 |
| HDPG | 24.88347 | 121.5019 | 197 | 62 | 33 | 185 | 3,201 | 37 | 1,201 | 475 | 475 |
| TTL | 23.45230 | 120.6134 | 215 | 79 | 45 | 174 | 3,515 | 98 | 2,154 | 81 | 123 |
| NAJY | 24.43352 | 121.7602 | 216 | 62 | 35 | 174 | 2,860 | 61 | 1,328 | 271 | 296 |
| HLCN | 23.90623 | 121.4928 | 213 | 68 | 40 | 168 | 2,127 | 46 | 849 | 251 | 255 |
| NXIR | 23.40887 | 121.4503 | 227 | 71 | 43 | 163 | 2,147 | 57 | 938 | 211 | 211 |
| DFR | 23.10356 | 121.2724 | 206 | 75 | 48 | 155 | 2,017 | 68 | 1,026 | 166 | 166 |

**Table S2.**
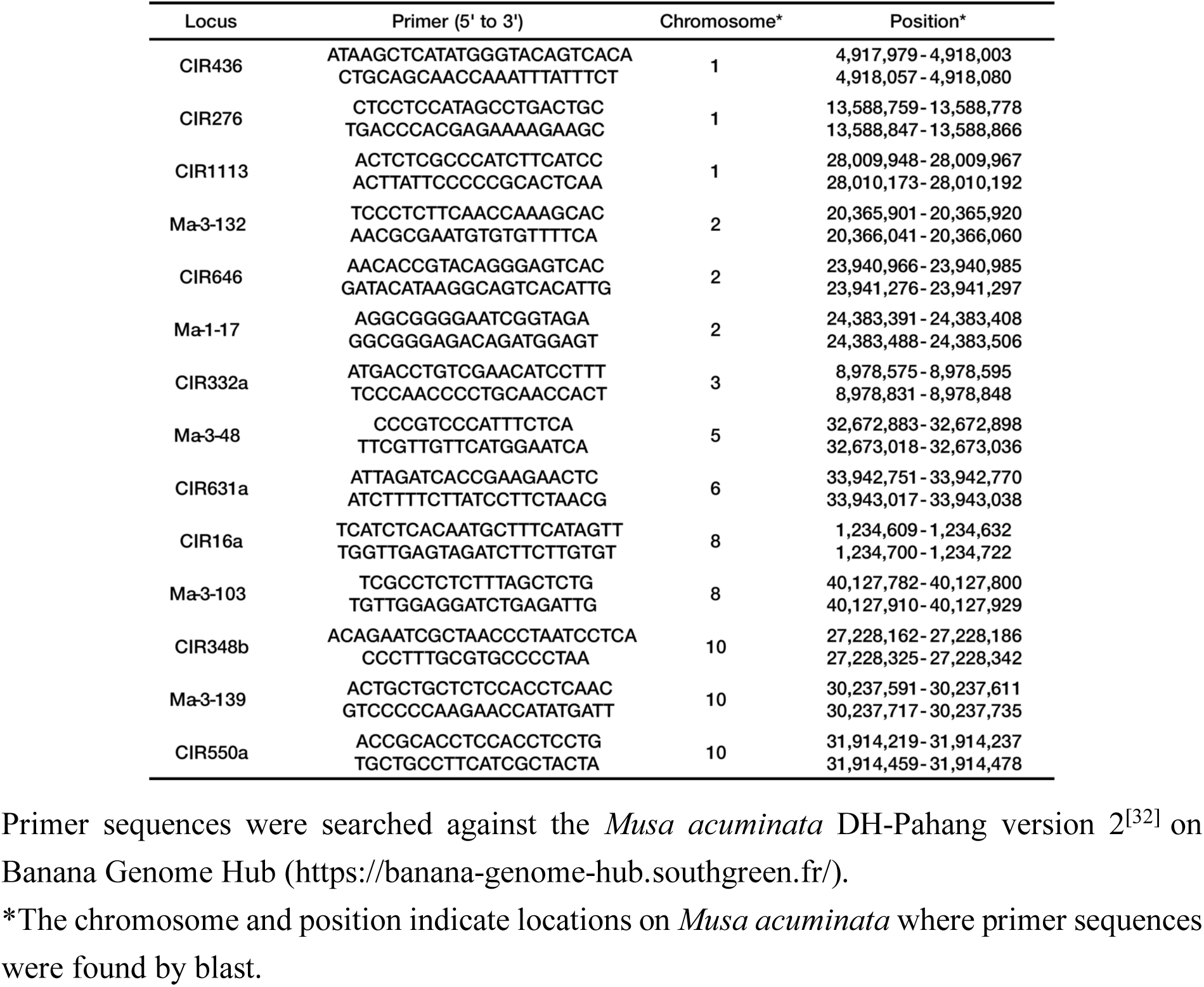
SSR primer information.

**Table S3.** Number of new-mutation (NM) or standing-variation (SV) SNPs with (adaptive) or without (non-adaptive) significant associations with bioclimatic variables

| Variable | Non-adaptive NM | Non-adaptive SV | Adaptive NM | Adaptive SV | *P* | Odds ratio |
| --- | --- | --- | --- | --- | --- | --- |
| BIO1 | 293,348 | 954,367 | 816 | 7,113 | 8.92e-169 | 2.68 |
| BIO2 | 293,274 | 953,198 | 890 | 8,282 | 8.43e-213 | 2.86 |
| BIO3 | 293,551 | 953,901 | 613 | 7,579 | 6.47e-256 | 3.80 |
| BIO7 | 293,644 | 953,981 | 520 | 7,499 | 1.32e-282 | 4.44 |
| BIO12 | 293,452 | 954,272 | 712 | 7,208 | 3.12e-203 | 3.11 |
| BIO15 | 293,168 | 951,447 | 996 | 10,033 | 2.33e-281 | 3.10 |
| BIO16 | 293,826 | 957,605 | 338 | 3,875 | 1.95e-123 | 3.52 |
| BIO17 | 293,461 | 954,969 | 703 | 6,511 | 1.61e-166 | 2.85 |
| BIO19 | 293,516 | 955,326 | 648 | 5,142 | 1.06e-107 | 2.44 |

Statistical significance from $\chi^2$ test is shown for each bioclimatic variable. Odds ratio is calculated as (adaptive SV / adaptive NM) / (non-adaptive SV / non-adaptive NM).

**Table S4.** Number of new-mutation (NM) or standing-variation (SV) SNPs with (adaptive) or without (non-adaptive) significant associations with bioclimatic variables (controlled for minor allele frequency)

| Variable | Non-adaptive NM | Non-adaptive SV | Adaptive NM | Adaptive SV | P | Odds ratio |
| --- | --- | --- | --- | --- | --- | --- |
| BIO1 | 44,920 | 260,113 | 238 | 2,573 | 1.27e-20 | 1.87 |
| BIO2 | 23,475 | 148,796 | 124 | 1,073 | 1.18e-03 | 1.37 |
| BIO3 | 39,432 | 234,613 | 182 | 2,689 | 2.21e-34 | 2.48 |
| BIO7 | 41,020 | 242,313 | 182 | 2,794 | 4.35e-38 | 2.60 |
| BIO12 | 37,266 | 224,455 | 220 | 2,514 | 3.36e-20 | 1.90 |
| BIO15 | 32,906 | 201,729 | 290 | 2,647 | 1.36e-10 | 1.49 |
| BIO16 | 33,685 | 206,816 | 113 | 1,282 | 2.88e-10 | 1.85 |
| BIO17 | 12,258 | 72,175 | 64 | 614 | 2.26e-04 | 1.60 |
| BIO19 | 11,237 | 66,547 | 41 | 362 | 1.81e-02 | 1.50 |

The number of four sets of SNPs whose minor allele frequency (MAF) ranges from the first quantile of adaptive MAF to the third quantile of non-adaptive MAF separately for each bioclimatic variable is shown. Statistical significance from $\chi^2$ test is shown for each bioclimatic variable. Odds ratio is calculated as (adaptive SV / adaptive NM) / (non-adaptive SV / non-adaptive NM).

**Table S5.** Number of currently adaptive new-mutation (NM) or standing-variation (SV) SNPs that remain (retention) or lose (disruption) significant associations with environments under all future climate-change scenarios

| Variable | Disruption<br>NM | Disruption<br>SV | Retention<br>NM | Retention<br>SV | *P* | Odds ratio |
| --- | --- | --- | --- | --- | --- | --- |
| BIO1 | 92 | 635 | 127 | 1,176 | 5.11e-02 | 1.30 |
| BIO2 | 34 | 374 | 348 | 2,010 | 6.85e-04 | 0.53 |
| BIO3 | 21 | 269 | 119 | 2,277 | 1.32e-01 | 1.50 |
| BIO7 | 29 | 251 | 128 | 2,393 | 4.53e-04 | 2.20 |
| BIO12 | 65 | 481 | 136 | 1,509 | 1.37e-02 | 1.50 |
| BIO15 | 37 | 365 | 437 | 2,982 | 4.79e-02 | 0.69 |
| BIO16 | 26 | 289 | 25 | 237 | 6.93e-01 | 0.85 |
| BIO17 | 37 | 354 | 159 | 1,182 | 2.21e-01 | 0.78 |
| BIO19 | 28 | 235 | 45 | 480 | 4.14e-01 | 1.30 |

Statistical significance from $\chi^2$ test is shown for each bioclimatic variable. Odds ratio is calculated as (retention SV / retention NM) / (disruption SV / disruption NM).

